# De novo protein structure prediction by incremental inter-residue geometries prediction and model quality assessment using deep learning

**DOI:** 10.1101/2022.01.11.475831

**Authors:** Jun Liu, Guang-Xing He, Kai-Long Zhao, Gui-Jun Zhang

## Abstract

**Motivation:** The successful application of deep learning has promoted progress in protein model quality assessment. How to use model quality assessment to further improve the accuracy of protein structure prediction, especially not reliant on the existing templates, is helpful for unraveling the folding mechanism. Here, we investigate whether model quality assessment can be introduced into structure prediction to form a closed-loop feedback, and iteratively improve the accuracy of de novo protein structure prediction.

**Results:** In this study, we propose a de novo protein structure prediction method called RocketX. In RocketX, a feedback mechanism is constructed through the geometric constraint prediction network GeomNet, the structural simulation module, and the model quality evaluation network EmaNet. In GeomNet, the co-evolutionary features extracted from MSA that search from the sequence databases are sent to an improved residual neural network to predict the inter-residue geometric constraints. The structure model is folded based on the predicted geometric constraints. In EmaNet, the 1D and 2D features are extracted from the folded model and sent to the deep residual neural network to estimate the inter-residue distance deviation and per-residue lDDT of the model, which will be fed back to GeomNet as dynamic features to correct the geometries prediction and progressively improve model accuracy. RocketX is tested on 483 benchmark proteins and 20 FM targets of CASP14. Experimental results show that the closed-loop feedback mechanism significantly contributes to the performance of RocketX, and the prediction accuracy of RocketX outperforms that of the state-of-the-art methods trRosetta (without templates) and RaptorX. In addition, the blind test results on CAMEO show that although no template is used, the prediction accuracy of RocketX on medium and hard targets is comparable to the advanced methods that integrate templates.

**Availability:** The RocketX web server are freely available at http://zhanglab-bioinf.com/RocketX.

## 1 Introduction

The accuracy of de novo protein structure prediction that does not rely on templates is limited for a long period of time due to the difficulty in designing accurate force field and efficient sampling algorithm (Rohl et al., 2004; Xu and Zhang, 2012; Kosciolek and Jones, 2014; Moult et al., 2018; Zhou et al., 2019; Liu et al., 2020a). With the exponential increase in protein sequences, since the Xu group applied the deep residual neural network (ResNet) to inter-residue contact prediction (Wang et al., 2017, 2018), the development of deep learning has promoted the rapid improvement in the accuracy of de novo structure prediction (Senior et al., 2020; Greener et al., 2019; Yang et al., 2020; Mortuza et al., 2021; Zhao et al., 2021). In general, the structure prediction methods based on deep learning are mainly divided into two categories: geometric constraint assisted folding (Xu, 2019; Zheng et al., 2019; Zhou et al., 2020; Ju et al., 2021; Su et al., 2021) and end-to-end prediction (Jumper et al., 2021; Baek et al., 2021). In AlphaFold2 (Jumper et al., 2021), MSA and structure templates are used as input, and attention-based networks (Evoformer) are used to generate expressions of single-residues and residue-pairs, and then 3D atomic coordinates are generated through the structural module based on invariant point attention (IPA). The network is trained end-to-end with the frame aligned point error (FAPE) loss and a number of auxiliary losses, and it finally achieves accurate end-to-end structure prediction. In CASP14, the GDT-TS score (Zemla, 2003) of AlphaFold2 on about two-thirds of the targets is higher than 90, which proves that consistent and high-precision protein structure prediction is possible. The end-to-end prediction has made a breakthrough, but the interpretability remains to be further studied (Skolnick et al., 2021). The geometric constraint assisted folding methods may be helpful for studying the multi-state of protein structure and revealing the folding mechanism.

Geometric constraint assisted folding generally includes two steps: geometric constraint prediction and structure folding. In geometric constraint prediction, a neural network model is usually trained to learn co-evolution relationships from the MSA of the query sequence to predict inter-residue contacts, distances and orientations. RaptorX introduces ResNet to modeling sequence features and pair features to predict inter-residue contacts (Wang et al., 2017; Xu, 2019). DeepMetaPSICOV (Jones et al., 2018) uses sequence profile, predicted secondary structure, solvent accessibility, and other features used in MetaPSICOV (Jones et al., 2015) as input to ResNet to predict inter-residue contacts. ResPRE (Li et al., 2019) uses the inverse covariance matrix of MSA to predict residue-level contact through ResNet. AmoebaContact (Mao et al., 2020) adopts a set of network architectures that are found as optimal for contact prediction through automatic searching, and it predicts the residue contacts at a series of cutoffs. In CASP13, the neural network has been extended to predict the inter-residue distance distribution (Senior et al., 2019; Xu and Wang, 2019; Hou et al., 2019; Kandathil et al., 2019; Kryshtafovych et al., 2019), which conveys more geometry information on the structure than contact. Furthermore, trRosetta (Yang et al., 2020; Du et al., 2021) introduces inter-residue orientations to characterize the geometric constraints in more detail. The predicted geometric constraints are converted into energy potential to guide structure folding through fragment assembly (Zhou et al., 2019; Liu et al., 2022; Mortuza et al., 2021; Xia et al., 2021) or energy minimization (Senior et al., 2020; Yang et al., 2020; Xu et al., 2021; Wang et al., 2021).

Model quality assessment methods have been proposed to evaluate the accuracy of the predicted models (Alapati and Bhattacharya, 2018; Cheng et al., 2019; Studer et al., 2020; Kwon et al., 2021; Ye et al., 2021). The traditional single-model quality assessment method uses various combinations of features and employ different machine-learning approaches for estimating the quality of a protein model (Wang et al., 2009; Uziela et al., 2016). In recent CASPs, a growing number of approaches have used deep learning and achieved impressive performance (Won et al., 2019; Huang et al., 2010; Baldassarre et al., 2021; McGuffin et al., 2021). ProQ3D (Uziela et al., 2017) shows that using exactly the same input as ProQ3 (Uziela et al., 2016) and only replacing the support vector machine with a deep neural network can improve the evaluation performance. Ornate (Pagès et al., 2019) constructs a residue voxelization feature that is rotationally and translationally invariant, and it predicts the local (residue-wise) and the global model quality through a deep 3D convolutional neural networks (CNN). 3DCNN (Derevyanko et al., 2018) and DeepAccNet (Hiranuma et al., 2021) also take advantage of 3D CNN to achieve advanced performance. Qdeep (Shuvo et al., 2020) integrates distance information through stack depth ResNets to evaluate model quality. Our recently proposed DeepUMQA (Guo et al., 2021) uses residue-level ultrafast shape recognition (USR) feature to describe the topological relationship between residues and the overall structure, and it combines other features to evaluate the quality of the model through CNN. DeepUMQA achieves top performance in CAMEO blind test. Model quality assessment is usually used as the last step to evaluate the quality of the model. Introducing it into the structure prediction to improve the model is worthy of attention.

In this work, we propose a de novo protein structure prediction method RocketX, in which a closed-loop feedback mechanism is constructed by inter-residue geometries prediction (GeomNet), structure simulation, and model quality assessment (EmaNet) to progressively improve the accuracy of predicted model. The structural model folded based on the geometric constraint predicted by GeomNet is evaluated by EmaNet, and the inter-residue distance deviation and per-residue lDDT estimated by EmaNet are fed back to GeomNet as the dynamic features to correct the geometric constraints prediction and progressively improve the structure model. Experimental results show that the closed-loop feedback mechanism can significantly improve the accuracy of geometric constraint prediction and the final model.

## 2 Methods

The pipeline of RocketX is shown in Figure 1, which consists of iterative three modules: inter-residue geometries prediction, structure simulation, and model quality assessment. For the query sequence, the MSA is generated by searched against the sequences databases using HHblits (Steinegger et al., 2019b). The features extracted from MSA and the dynamic features fed back from model assessment network are used as input to an improved deep residual neural network (GeomNet) to predict the inter-residue geometries. Then, the predicted geometric constraints are converted into smooth energy potentials to guide the structure folding by direct energy minimization and full-atom relaxation under the framework of PyRosetta (Chaudhury et al., 2010). The structure features extracted from the folded model are fed into a deep convolutional neural network (EmaNet) to estimate the inter-residue deviation and per-residue lDDT of the model, which will be fed back to the GeomNet as the dynamic features to correct the geometries prediction and progressively improve the structure model. The final model is generated after three iterations.

**Fig. 1.**
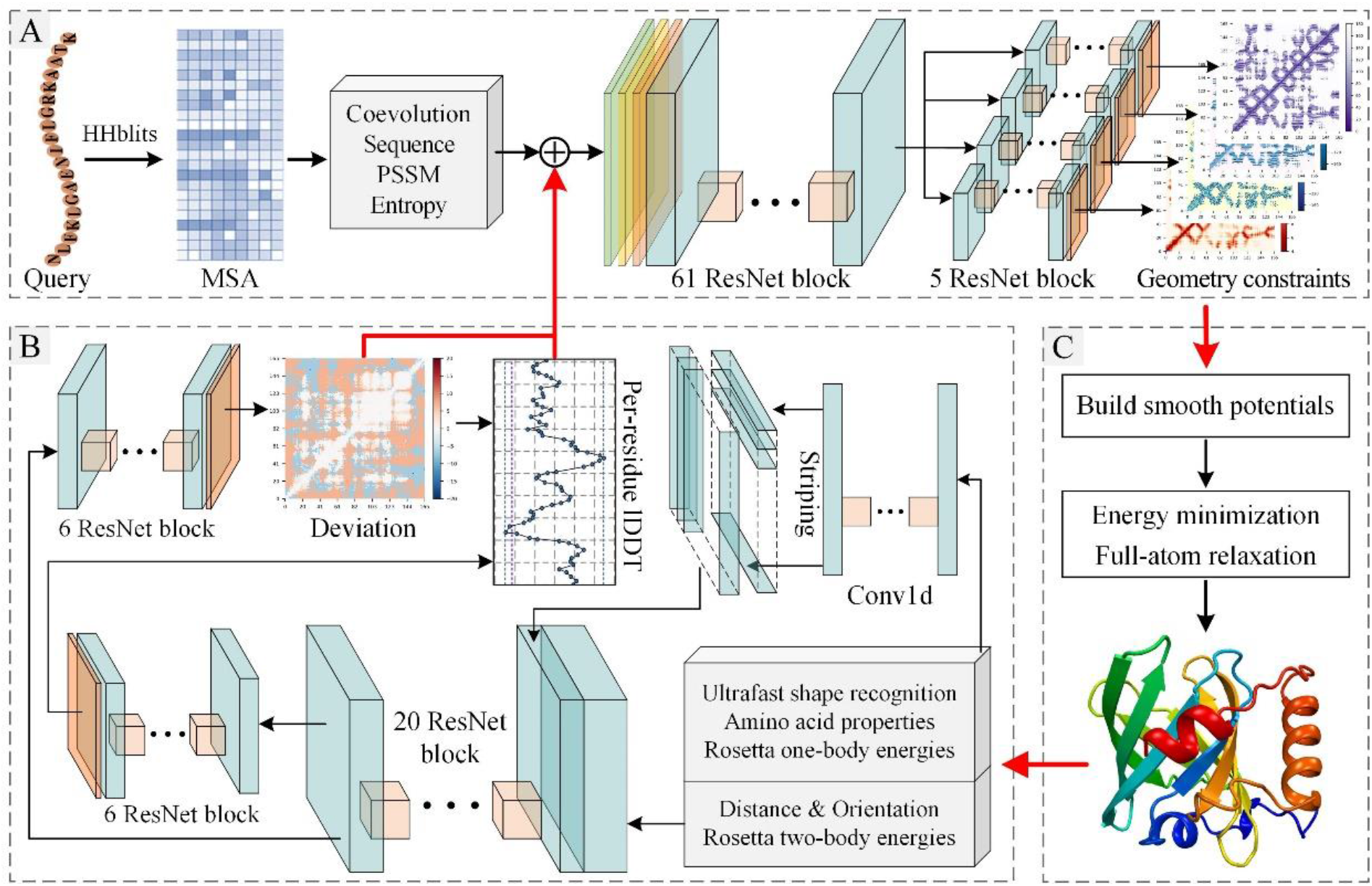
RocketX pipeline. (A) Geometric constraints prediction. The co-evolutionary features extracted from the MSA of query sequence and the dynamic features fed back by model quality assessment neural network are fed into the deep residual neural network to predict the inter-residue geometric constraints. (B) Structure folding. The predicted geometric constraints are converted into smooth potentials, and the structure model is generated by direct energy minimization and full-atom relaxation. (C) The predicted structure model is sent to the model assessment neural network to predict the inter-residue distance deviation and per-residue lDDT that will be fed back to (A) to correct the inter-residue geometries prediction and fold the structure model.

### 2.1 Benchmark datasets

In this work, the dataset of DeepAccNet (Hiranuma et al., 2021) is used to train our model assessment network. It contains 7800 proteins (95% for training, 5% for validation) collected from the PICSCES server (deposited by May 1, 2018) with residues from 50 to 300. Each protein has approximately 150 decoy structures generated through comparative modeling, native structure perturbation, and deep learning guided folding. The benchmark dataset of geometries prediction network was constructed from SCOPe 2.07 (Fox et al., 2014) (deposited by November 30, 2018). To train the geometric constraints prediction neural network, the sequences with the sequence similarity of more than 40% with the dataset of model evaluation network are eliminated by CD-HIT-2D (Jing et al., 2020), and 166,025 sequences are retained. Then, the remaining sequences are limited to 50 to 500 residues and clustered by CD-HIT (Fu et al., 2012) with 30% sequence similarity to generate 9,091 representatives. A total of 8608 sequences (95% for training, 5% for validation) are randomly selected to train the neural network, and the remaining 483 sequences are used as the benchmark test set of the overall method. In addition, 20 FM targets of CASP14 are used to further test the performance of RocketX. RocketX also participate in the blind test of CAMEO.

### 2.2 Geometric constraints prediction

#### 2.2.1 MSAs collection and features extraction

For proteins in training and test set, MSA is generated by iteratively search against UniRef30 (Mirdita et al., 2017) and BFD (Steinegger et al., 2019a) sequence databases using HHblits (Steinegger et al., 2019b). The gradually relaxed e-value cutoffs (1e^−30^, 1e^−10^, 1e^−6^, 1e^−3^) are used until the MSA has at least 2,000 sequences with 75% coverage or at least 5,000 sequences with 50% coverage. The input of the geometric constraint prediction neural network includes the conventional features extracted from MSA: 1) inverse of covariance matrix (441 features × L × L) and average product correction (1 features × L), 2) position specific scoring matrix (21 features × L), 3) residue position entropy (1 features × L) and 4) one-hot-encoding of the query sequence (20 features × L); and the dynamic features that are fed back from the model evaluation neural network: 5) inter-residue distance deviation (15 features × L), and 6) per-residue lDDT (1 features × L). Features defined on single residues are converted into 2D maps by horizontally and vertically striping, and then, they are sent to the network together with 2D features. Only the features from MSA are used in the first round of geometric constraints prediction.

#### 2.2.2 Network architecture

Figure 1A shows a high-level overview of the geometric constraints prediction neural network, and the detailed network architecture can be found in Supplementary Figure S1. The network takes the abovementioned features as input to predict the inter-residue distance and orientation distributions through a trunk residual block and four branch residual blocks, respectively. The definitions of inter-residue distance and orientation are same as trRosetta (Yang et al., 2020). The distance range (from 2 to 20Å) is divided into 36 equidistant bins in step of 0.5Å plus one bin indicating that the residues are not in contact. Similarly, ω, θ dihedrals, and φ angle are divided into 24, 24, and 12 bins, respectively, in step of 15Å (plus one no-contact bin). The 2D convolutional layer is first used to reduce the number of channels of the input tensor to 64, and then, 61 basic residual blocks are applied. Each residual block consists of two 2D convolutional layers (with dilation size cycle through 1, 2, 4, 8, and the last is 1), two instance normalization layers, two ELU activation layers, and a dropout layer with 15% drop probability. The network branches into four independent paths after the trunk block to learn the potential characteristics of different geometric constraints. Each branch consists of 5 basic residual blocks (the same as the basic residual blocks in the trunk), a softmax activation layer, and a 2D convolutional layer (convert the number of output channels to 37, 25, 25, and 13).

#### 2.2.2 Model training

We train two network models for geometric constraints prediction using the same parameters. The first one is used for the first round prediction, and the input tensor is only the features extracted from MSA (L × L× 526). The other is used for subsequent prediction, and the input includes the features extracted from MSA and the features fed back from the model assessment network (L × L× 543). During training, AdamW optimizer with a "triangular" cycle learning rate (base_lr=1e^−4^, max_lr—5e^−4^, step_size_up=10, step_size_down=10) is used. The categorical cross-entropy is used to measure the loss for the four kinds of geometries. The total loss is the sum of the four individual losses with same weight. A total of 200 epoches of training are performed, and each epoch traversed the entire training set. The five models with the lowest loss on the validation set are saved, and the average results of the five models as used as the final prediction. The network takes about 10 days to train on one NVIDIA TITAN RTX GPU.

### 2.3 Structure folding

In this study, we build protein tertiary structure model from the predicted inter-residue geometric constraints. First, the predicted inter-residue distance and orientation distributions are converted into energy potentials using the DFIRE (Zhou and Zhou, 2002) paradigm. The energy potentials are smoothed by the spline function in Rosetta and the four Rosetta basic energy terms are combined to guide the accurate generation of the structure model. Then, *MinMover* in PyRosetta (Chaudhury et al., 2010) is used to search for the coarse-grained tertiary structure model with the minimal potential. Finally, the coarse-grained model is refined into full-atom model by executing *FastRelax*.

### 2.4 Model assessment network

#### 2.2.1 Feature collection

The input features of the model assessment neural network are purely extracted from the predicted structure model, including 1D features (i.e., per-residue USR, amino acid properties, secondary structure, bond lengths and angles, backbone dihedral angles, and Rosetta one-body energy terms), and 2D features (i.e., inter-residue multi-distances and orientations, hydrogen bond maps, Euler angles of residue pairs, sequence separation, and Rosetta two-body energy terms). The detailed feature descriptions are shown in Supplementary Table S1. Specially, we have proposed residue-level USR feature to characterize the relationship between individual residues and topological structure (Guo et al., 2021). Briefly, for the current residue *ri*, the residue *ri*’ that farthest from *ri* and the residue *ri*’’ that farthest from *ri*’ are first selected, and then the first moments of distances from all residues to the three residues are calculated as residue-level topological feature. Amino acid properties include one-hot encoding of amino acid, Blosum62 scores (Henikoff et al., 1992), and per amino-acid features from Meiler (Henikoff et al., 1992). The secondary structure is calculated by DSSP (Kabsch and Sander, 1983) and expressed as one-hot encoding of three states. The distances between different atoms of inter-residue are used to describe the geometric characteristics of the structure model, including Cβ to Cβ (Cα for GLY) distance map, Cα to tip atom distance map, tip atom to Cα distance map, and tip atom to tip atom distance map. The definition of tip atom for each residue are listed in Supplementary Table S2.

#### 2.2.1 Network architecture

The high-level overview of the model assessment neural network is shown in Figure 1C, and the detailed network architecture can be found in Supplementary Figure S2. The input of the network consists of 73 1D features and 33 2D features. The 1D features first pass through the 1D convolutional layer into 64 channels. It is then converted into a 2D tensor by horizontally and vertically striping, and combined with the input 2D features into a 161 channel 2D tensor. The 2D tensor is transformed into a standard 128 channel through the 2D convolution layer, instance normalization layer, and ELU activation layer. This procedure is followed by the trunk ResNet operations of the network. The trunk ResNet operations consist of a 2D convolutional layer, 20 basic residual blocks, and an ELU activation layer. Each basic residual block contains three 2D convolutional layers, three instance normalization layers, and three ELU activation layers. The kernel size of the three convolutional layers are 1×1, 3×3, and 1×1, respectively. The dilation size of the second layer is cycle through [1, 2, 4, 8]. The first and third convolutional layers are used to reduce and restore the number of channels of the tensor (128→64→128). The network is then branches into two independent paths after the trunk ResNet. The branch ResNet operations is similar to the trunk ResNet, except that the instance normalization layer is removed. The ResNet operation of each branch is followed by a 2D convolutional layer and a softmax activation layer or sigmoid activation layer for predicting the inter-residual distance deviation or the contact map with a threshold of 15° A. Then, the per-residue lDDT is calculated according to the output of the two branches, and the inter-residual distance deviation and per-residue lDDT are used as the output of the model assessment network.

#### 2.2.1 Model training

In order to reflect the quality of the model in detail so as to feed it back to the geometric constraint prediction network for correction. We use signed inter-residue distance deviation and per-residue lDDT as the output of the model assessment network. The inter-residue distance deviation is defined as the difference between the inter-residue distance (Cβ atom, Cα atom for GLY) of the structure model and the native structure. It is divided into 15 bins with the boundaries of ±0.5, ±1, ±2, ±4, ±10, ±15, and ±20Å. lDDT (Mariani et al., 2013) is a superposition-free score that evaluates local distance differences of all atoms in a model. In this work, the lDDT of each residue is calculated by the predicted inter-residue distance deviation and the predicted contact map. For residue *i*, the lDDT is defined as follows:

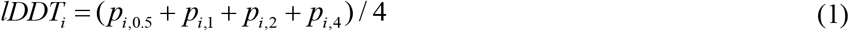

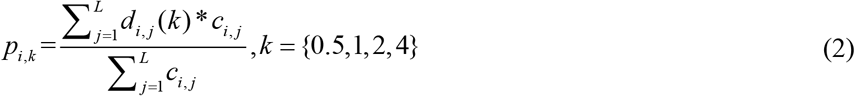

where *p_i,k_* is the predicted lDDT of residue i under the threshold of k; *d_i,j_* (*k*) is the probability that the distance deviation of residues *i* and *j* is within ±*k*Å; *c_i,j_* is the probability that residues *i* and *j* are contact (within 15Å); *L* is the number of residues. The final lDDT score of the residue is the average of the lDDT computed using the threshold of 0.5, 1, 2, and 4Å.

During the training, Adam optimizer with a learning rate of 0.0005 and a decay rate of 0.98 per epoch is used. The categorical cross-entropy is used to measure the loss of inter-residue distance deviation and contact, and the mean squared is used to measure the loss of per-residue lDDT. The total loss is the sum of the three losses with weights of 1, 0.25, and 10. The weights are tuned to allows the deviation to dominate because it can provide richer information for subsequent optimization. Contacts does not participate in subsequent optimization as feedback, so a smaller weight is assigned. At each step of training, a single decoy is randomly selected from the decoy set of a training protein. A total of 200 epochs are trained, and all training proteins were randomly traversed in each epoch. The five models with the lowest loss on the validation set are saved, and the average results of the 5 models as used as the final prediction. The network takes about 8 days to train on one NVIDIA TITAN RTX GPU.

## 3 Result and discussion

### 3.1 Accuracy of predicted inter-residue geometries

RocketX is used to evaluate the inter-residue geometries for the 504 benchmark test proteins and is compared with trRosetta. For fair comparison, we use the distance evaluation server DISTEVAL (Adhikari, 2021) to evaluate the precision of the predicted distances. The comparison results are summarized in Table 1, and the detailed results are shown in Supplementary Table S3.

**Table 1.** Precision of the predicted inter-residue geometries of RocketX and trRosetta on the benchmark testset, which is evaluated from two levels of contact and distance. *Note*: The second and third columns are the precision of top L predicted contacts with sequence separation (sep) ≥12 and ≥24, respectively. The fourth and fifth columns are the mean absolute error (MAE) and root mean squared error (RMSE) of predicted and real medium-long-range distances. The six column is the Cβ-lDDT calculated according to the predicted and real distances.

| Method | Contact level |  | Distance level |  |  |
| --- | --- | --- | --- | --- | --- |
| | $sep \geq 12$ | $sep \geq 24$ | MAE | RMSE | $C_\beta$-IDDT |
| RocketX | 0.764 | 0.667 | 1.09 | 1.50 | 0.677 |
| trRosetta | 0.749 | 0.650 | 1.15 | 1.57 | 0.623 |

At the contact level, the sum of the probabilities of the intervals with a distance below 8Å in the distance distribution serves as the predicted contact probability. The contact precision of RocketX’s medium-long-range (*sep*≥12) and long-range (*sep*≥24) are 0.764 and 0.667, respectively, which are 2.00% and 2.62% higher than those of trRosetta. At the distance level, the mean absolute error (MAE) and root mean squared error (RMSE) of predicted and real medium-long-range distances are used to measure the distance precision. The Cβ-lDDT calculated from the predicted and real distance is also used. The MAE and RMSE of RocketX are 1.08Å and 1.50Å, respectively, which are 5.56% and 4.46%lower than those of trRosetta. The Cβ-lDDT of RocketX is 0.677, which is 8.65% higher than that of trRosetta. Figure 2(A) intuitively reflect the comparison of the top L medium-long-range contact precision with sequence separation≥12 of RocketX and trRosetta. RocketX achieves higher contact precision on 334 of 483 proteins, which account for 69.2%of the total. The head-to-head comparison of the C_*β*_-lDDT calculated from the predicted inter-residue distances of RocketX and trRosetta is shown in Figure 2(B). RocketX achieves higher Cβ-lDDT score on 414 proteins, which account for 85.7% of the total. The intuitively comparison of top *L* long-range contact precision, the C*β*-lDDT with sequence separation ≥6, the MAE, and the RMSE can be found in Supplementary Figure S3.

**Fig. 2.**
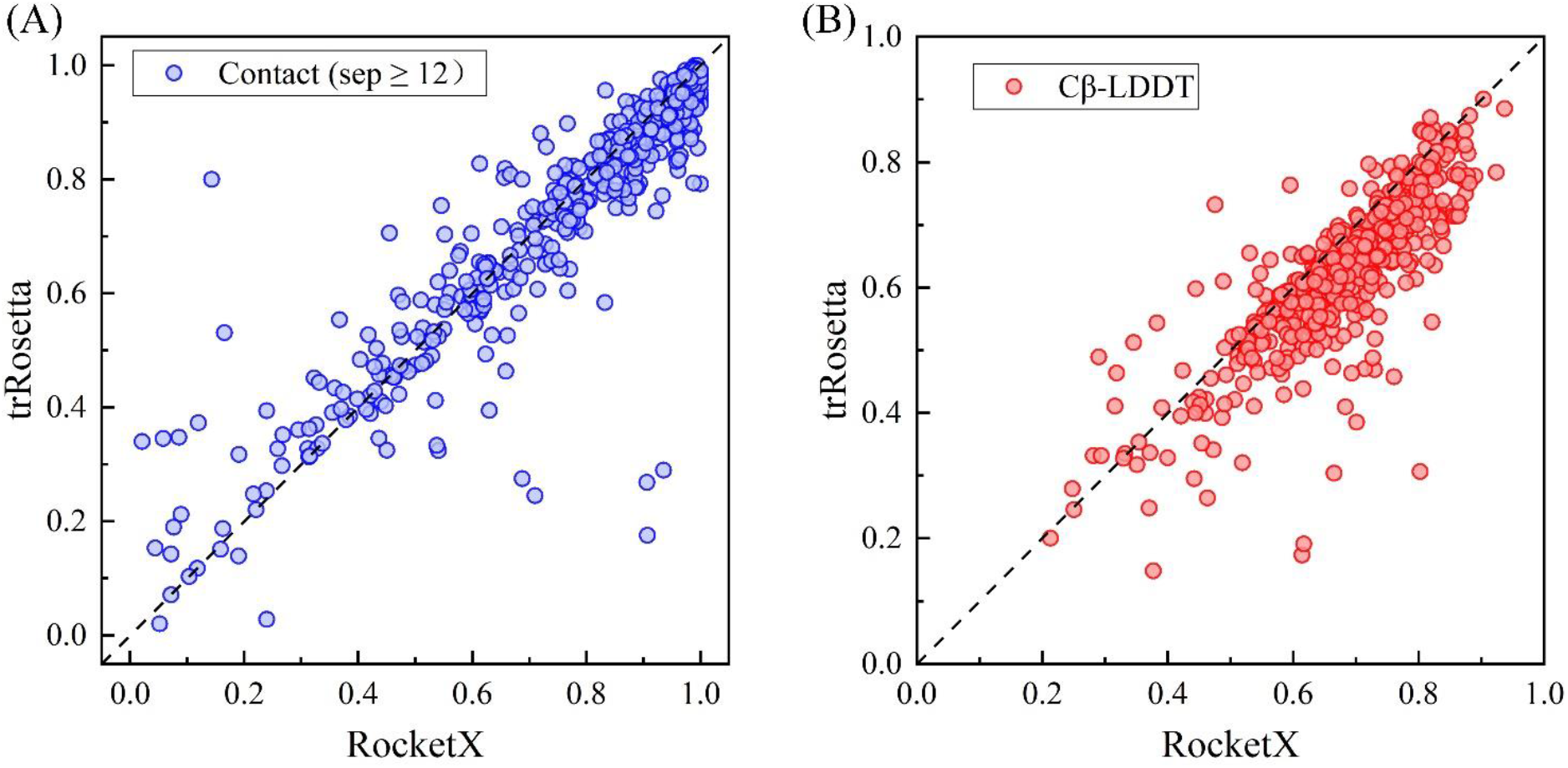
(A) Head-to-head comparison of the contact precision with sequence separation ≥12 predicted by RocketX and trRosetta. (B) Head-to-head comparison of the Cβ-lDDT calculated from the predicted inter-residue distances of RocketX and trRosetta.

### 3.2 Comparison with the state-of-the-art methods

To test the performance of the overall method, we also evaluate the accuracy of the structure model predicted by RocketX and compare it with the state-of-the-art methods trRosetta and RaptorX. The results of trRosetta are predicted from its official server with the "*Do not use templates*" option selected (http://yanglab.nankai.edu.cn/trRosetta/). The results of RaptorX are predicted from its official server (http://raptorx.uchicago.edu/ContactMap/). The prediction results on the benchmark testset are summarized in Table 2, and the detailed results of each protein are listed in Supplementary Table S4.

**Table 2.** Results of RocketX, trRosetta, and RaptorX on the benchmark test set. The last column is the result of the Wilcoxon signed-rank test calculated in accordance with TM-score.

| Method | RMSD | TM-score | TM-score $\geq 0.5$ | TM-score $\geq 0.8$ | TM-score $\geq 0.9$ | $p$-value |
| --- | --- | --- | --- | --- | --- | --- |
| RocketX | 4.92 | 0.774 | 457 | 263 | 81 | NA |
| trRosetta | 5.17 | 0.751 | 455 | 216 | 37 | 5.54E-14 |
| RaptorX | 5.88 | 0.714 | 429 | 188 | 25 | 4.96E-21 |

The average RMSD of RocketX is 4.92, which is 4.84% and 16.33%lower than those of trRosetta and RaptorX, respectively. The average TM-score (Xu and Zhang, 2010) of RocketX is 0.774, which is 3.06% and 8.40% higher than those of trRosetta and RaptorX, respectively. RocketX correctly folds (i.e. TM-score≥0.5) 457 out of 483 proteins, which account for 94.6% of the total and are more than those of trRosetta and RaptorX. In addition, RocketX predicts a model with TM-score≥0.8 on 263 proteins, which is 47 and 75 more than those of trRosetta and RaptorX, respectively. The number of models with TM-score≥0.9 of RocketX is 81, which is 2.19 and 3.24 times those of trRosetta and RaptorX, respectively. The result of Wilcoxon signed-rank test (Corder and Foreman, 2009) calculated in accordance with TM-score shows that RocketX is significantly better than trRosetta and RaptorX in the benchmark test set. Figure 3 more intuitively reflects the comparison of the TM-score of RocketX, trRosetta, and RaptorX. Compared with trRosetta, RocketX achieves a higher TM-score on 326 proteins, which account for 67.5% of the total; and the TM-score of RocketX is increased by more than 0.1 over trRosetta on 45 proteins. Compared with RaptorX, RocketX achieves a higher TM-score on 314 proteins, which account for 65.0% of the total; and the TM-score of RocketX is increased by more than 0.1 over trRosetta on 115 proteins. Linear fitting shows that RocketX has more advantages in generating high-precision models than trRosetta and RaptorX. Figure 3(C) shows the number of models of RocketX, trRosetta, and RaptorX in each TM-score interval (with a step of 0.05). In the intervals of TM-score<0.7, RaptorX has the most models; in the intervals of from 0.7 to 0.8, trRosetta has the most models; in the interval of 0.8~0.85, trRosetta and RocketX have the same number of models, which is more than that of RaptorX; in the intervals of from 0.85 to 1.0, RocketX has the most models. Figure 3(D) shows the boxplot of the TM-score of the models predicted by RocketX, trRosetta and RaptorX. The Q1 (first quartile), Q2 (second quartile or median), and Q3 (third quartile) of RocketX are 0.698, 0.814 and 0.878, those of trRosetta are 0.682, 0.780 and 0.853, and those of RaptorX are 0.615, 0.753, and 0.843.

**Fig. 3.**
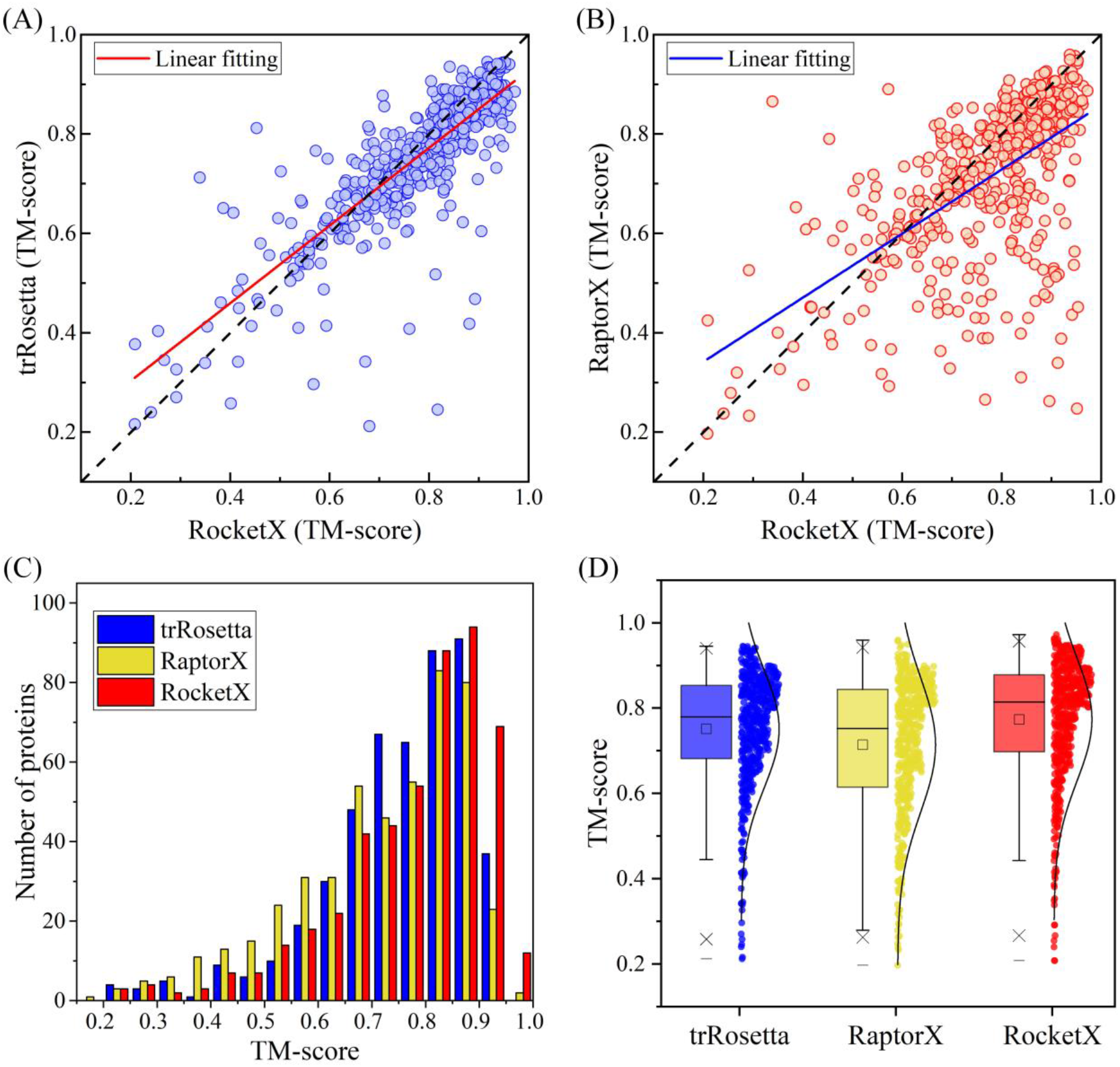
(A) Head-to-head comparison of TM-score between RocketX and trRosetta. (B) Head-to-head comparison of TM-score between RocketX and RaptorX. (C) TM-score distribution of structure models predicted by RocketX, trRosetta, and RaptorX in each TM-score intervals (with step of 0.05). (D) TM-score boxplot of the models predicted by RocketX, trRosetta, and RaptorX.

### 3.3 Contribution of model assessment feedback

We design two ablation experiments to investigate the contribution of model assessment feedback. The first is RocketX^1^, which does not use model evaluation feedback, but only uses the features extracted from MSA to predict geometric constraints and fold structural models. The second is RocketX^2^ that uses one round of model assessment feedback. Note that RocketX uses two rounds of model assessment feedback. The prediction results of different version of RocketX on the benchmark testset are summarized in Table 3, and the detailed results of each protein are listed in Supplementary Table S4.

**Table 3.** Results of RocketX, RocketX^2^, and RocketX^1^ on the benchmark test set. *Note*: RocketX^1^ is a version of RocketX that does not use model assessment feedback. RocketX^2^ is a version of RocketX that uses one round of model assessment feedback. Note that RocketX uses two rounds of model assessment feedback.

| Method | RMSD | TM-score | TM-score$\geq 0.5$ | TM-score$\geq 0.8$ | TM-score$\geq 0.9$ | *p*-value |
| --- | --- | --- | --- | --- | --- | --- |
| RocketX | 4.92 | 0.774 | 457 | 263 | 81 | NA |
| RocketX<sup>2</sup> | 4.99 | 0.768 | 457 | 251 | 73 | 1.81E-29 |
| RocketX<sup>1</sup> | 5.12 | 0.758 | 456 | 233 | 55 | 2.71E-57 |

The average RMSD and TM-score of RocketX^1^ are 5.12 and 0.758, respectively. The average RMSD of RocketX^2^ is decreased by 2.54%, and the average TM-score is increased by 1.32% when one round model assessment feedback is considered. The average RMSD of RocketX is decreased by 3.91%, and the average TM-score is increased by 2.11% when two rounds of model assessment feedback are considered. The number of correctly folded models in RocketX^1^ is 456, and the number of RocketX^2^ and RocketX models has increased by one. RocketX^1^ obtained models with TM-score≥0.8 on 233 proteins. On this basis, RocketX^2^ has increased by 18, and RocketX has further increased by 12 compared with RocketX^2^. The number of models with TM-score≥0.9 of RocketX^1^ is 55, and those of RocketX^2^ and RocketX increase to 73 and 81, respectively. RocketX^2^ achieves higher TM-score than RocketX^1^ on 409 proteins, which account for 84.7% of the total. Furthermore, RocketX achieves higher TM-score than RocketX2 350 proteins, which account for 72.5% of the total. The Wilcoxon signed-rank test result show that the contribution of model assessment feedback is significant. Figure 4 more intuitively reflects the comparison of the TM-score of different versions of RocketX. Of particular note is the exciting increase in the number of TM-score≥0.95 models from 1 in the RocketX^1^ to 12 in the RocketX due to the addition of model evaluation feedback (Figure 4(A)). In the boxplot of Figure 4(B), Q1, Q2, and Q3 of the TM-score of RocketX^1^ are 0.680, 0.794, and 0.863, respectively, and those of RocketX^2^ are 0.691, 0.806, and 0.873. Figure 4(C) clearly shows the gain from model evaluation feedback on each benchmark test protein.

**Fig. 4.**
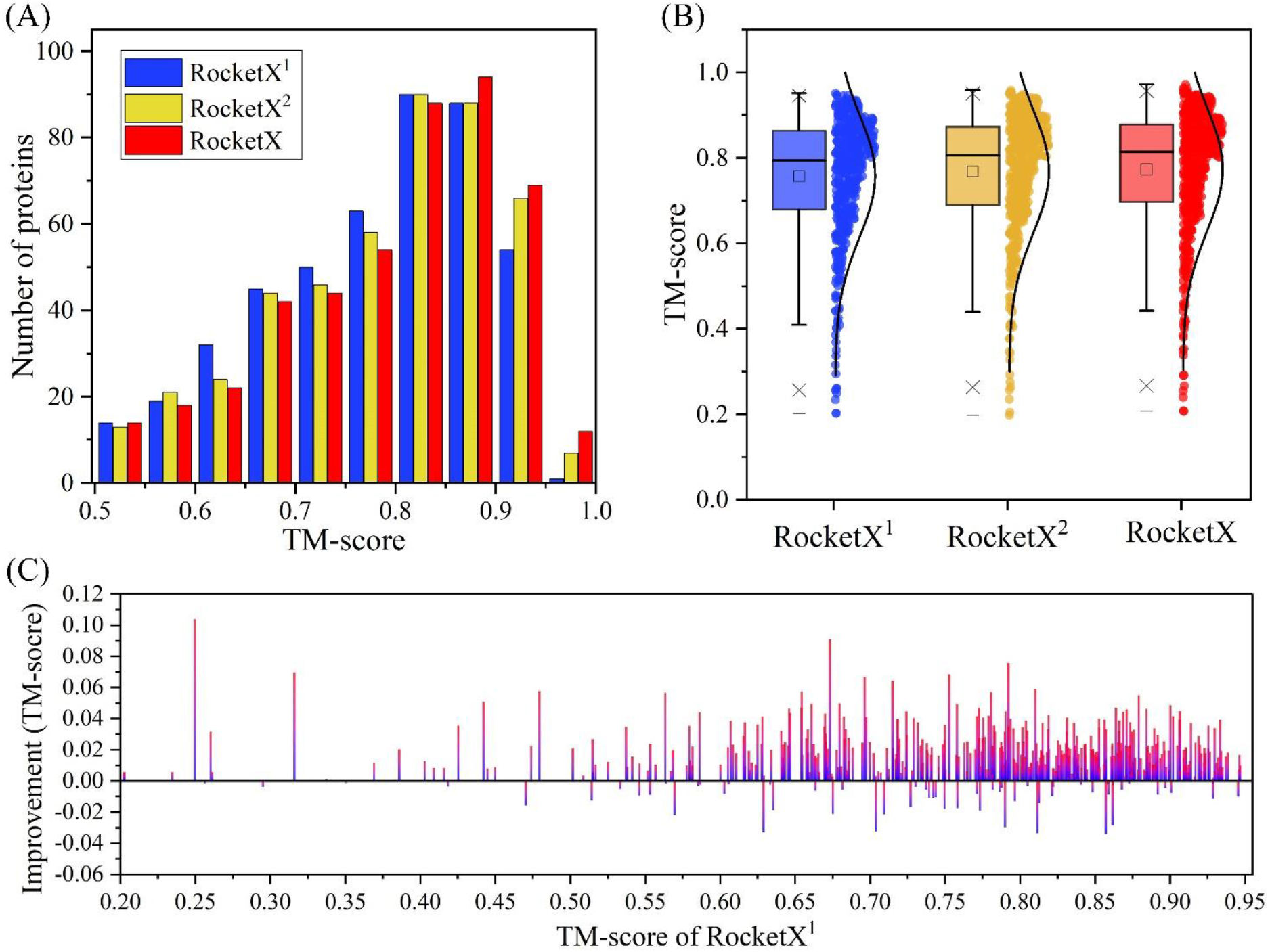
(A) Number of structure models of RocketX^1^, RocketX^2^, and RocketX in different TM-score intervals. (B) Boxplot of TM-score of the models predicted by RocketX^1^, RocketX^2^, and RocketX. (C) Accuracy improvement brought by model evaluation feedback on each benchmark protein. The horizontal axis is the accuracy before the model evaluation feedback, and the vertical axis is the change in accuracy after the model evaluation feedback is added.

### 3.4 Contribution of the network architecture of geometries prediction

We analyze the performance of geometric constraints prediction under two network architectures. The first is similar to trRosetta, which only uses a series of ResNet blocks on the trunk of the network and then uses 2D convolution to directly map into four prediction components, denoted as GeomNet^*^. The second is the one we used, that is, after the main ResNet operation, which is to use ResNet blocks on the four network branches after the trunk ResNet blocks to learn the potential properties of the different components, named GeomNet. We train the two networks with the same training set and parameters and compare them on the benchmark test set using the same MSA as input. The comparison results are summarized in Table 4, and the detailed results are listed in Supplementary Table S5. The precision of the geometric constraints predicted by GeomNet is better than that of GeomNet^*^ whether at the contact or the distance level. Furthermore, at the level of using predicted geometric constraints to build structure models, GeomNet outperforms GeomNet^*^ on all evaluation criteria. The significance test results show that using the geometric constraints predicted by GeomNet to build a model is significantly better than using those by GeomNet^*^. A more intuitive and detailed comparison can be found in Supplementary Figure S4. Experimental results show that using ResNet blocks in the branch network that predict different geometric constraints can achieve more accurate prediction than using only ResNet blocks in the trunk. We speculate that different inter-residue geometries have common and own characteristics. The trunk network can learn their common characteristics, and the branch network may learn their respective potential characteristics.

**Table 4.** Performances of the two network architecture of the geometry constraints prediction on the benchmark test set. *Note*: GeomNet^*^ is the version of network that only uses ResNet blocks on the trunk of the network. GeomNet is the network we used, in which the ResNet blocks are also used on the network branches after the trunk ResNet blocks. Columns 8 to 13 are the evaluation of the structural model constructed based on the predicted geometric constraints. #TM0.5, #TM0.8, and #TM0.9 are the number of proteins with TM-score≥0.5, TM-score≥0.8, TM-score≥0.9, respectively.

| Method | Contact level |  | Distance level |  |  | RMSD | TM-score | #TM0.5 | #TM0.8 | #TM0.9 | *p*-value |
| --- | --- | --- | --- | --- | --- | --- | --- | --- | --- | --- | --- |
| | *sep*≥12 | *sep*≥24 | MAE | RMSE | C$_{\beta}$-IDDT | | | | | | |
| GomNet | 0.743 | 0.643 | 1.15 | 1.57 | 0.616 | 5.12 | 0.758 | 456 | 233 | 55 | NA |
| GomNet* | 0.731 | 0.630 | 1.22 | 1.66 | 0.602 | 5.36 | 0.736 | 459 | 189 | 21 | 6.73E-26 |

### 3.5 Performances on CASP targets

We also tested the performance of RocketX on 20 FM targets of CASP14 targets and compared it with four state-of-the-art methods of server groups in CASP14, namely RaptorX, BAKER-ROSETTASERVER, MULTICOM-CLUSTER, and Yang_FM. The results of the compared methods are obtained from the CASP official website (http://predictioncenter.org). Figure 5 shows the TM-score of the first model predicted by each method on each target, and the detailed results are listed in Supplementary Table S6. The average TM-score of RocketX on the 20 targets is 14.11% higher than that of RaptorX (0.415), 15.51% higher than that of BAKER-ROSETTASERVER (0.410), 3.05%higher than that of MULTICOM-CLUSTER, and 0.64% higher than that of Yang_FM (0.470). The number of correct folding targets for RocketX is 7, and that of RaptorX, BAKER-ROSETTASERVER, MULTICOM-CLUSTER, and Yang_FM are 5, 6, 9, and 9 respectively. RocketX achieved the highest TM-scroe on 6 out of the 20 targets, and those of RaptorX, BAKER-ROSETTASERVER, MULTICOM-CLUSTER, and Yang_FM are 2, 10, 1, and 2, respectively.

**Fig. 5.**
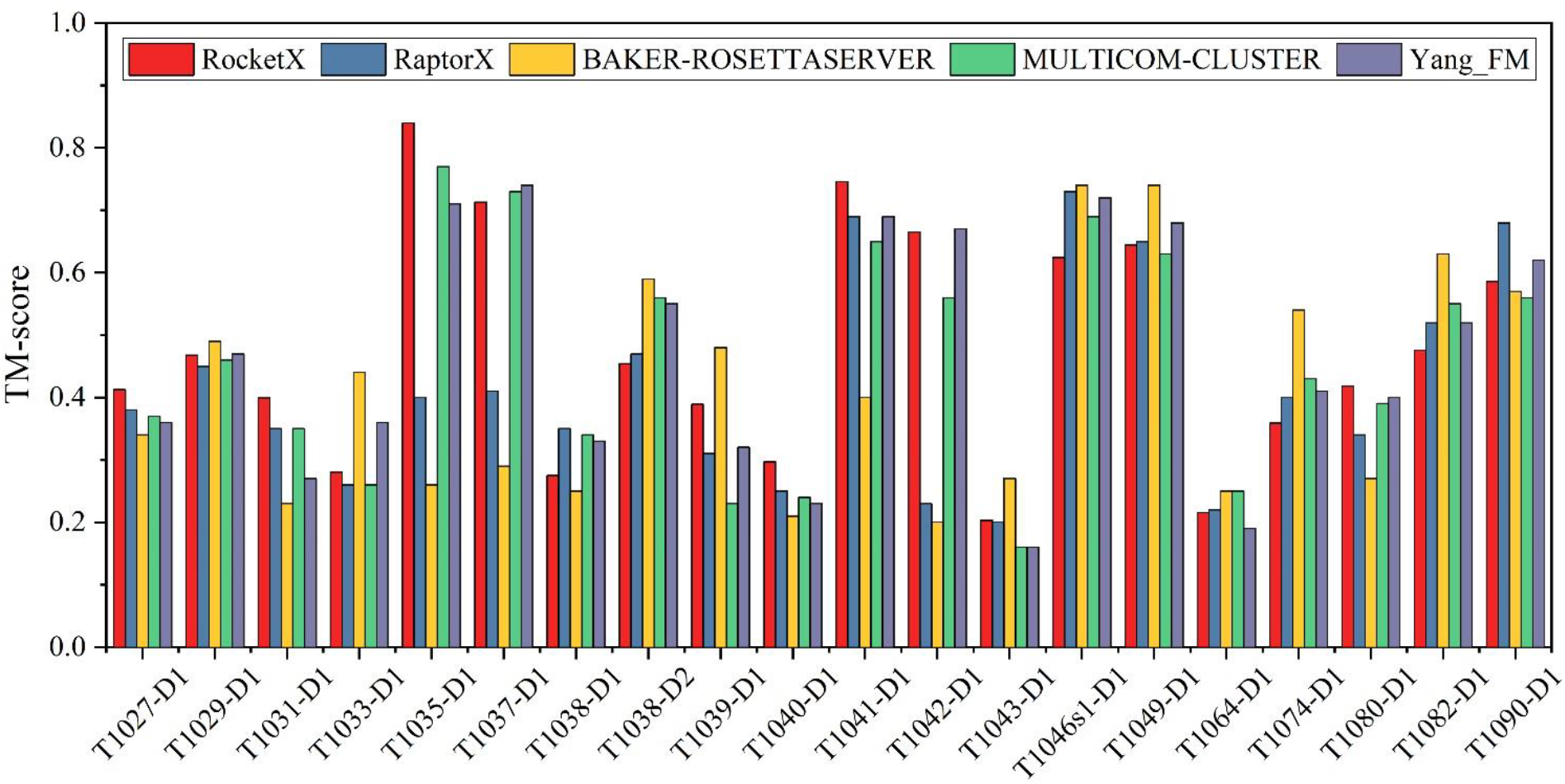
TM-score of the first model predicted by RocketX, RaptorX, BAKER-ROSETTASERVER, MULTICOM-CLUSTER, and Yang_FM on the 20 FM targets of CASP14.

### 3.5 Performances on CAMEO blind test

RocketX server also participate in the blind test of CAMEO (https://www.cameo3d.org/modeling) for 1 month (2021-12-04 to 2021-12-25). Figure 6 and Supplementary Figures S5 and S6 shows the the performance of RocketX and 10 servers in the CAMEO, namely, RoseTTAFold, Robetta, IntFold6-TS, IntFold5-TS, IntFold4-TS, IntFold3-TS, RaptorX, SWISS-MODEL, PRIMO, and Phyre2. The detailed server introduction can be found on the CAMEO official website (https://www.cameo3d.org/cameong_servers/3D/). Notably, all these methods use templates, while RocketX does not use any template information. On the 16 hard targets, the average TM-score and lDDT of RocketX are 0.45 and 43.58, respectively, which are lower than those of RoseTTAFold, Robetta and IntFold6-TS but higher than those of other methods, ranking 4th out of 11 methods.

**Fig. 6.**
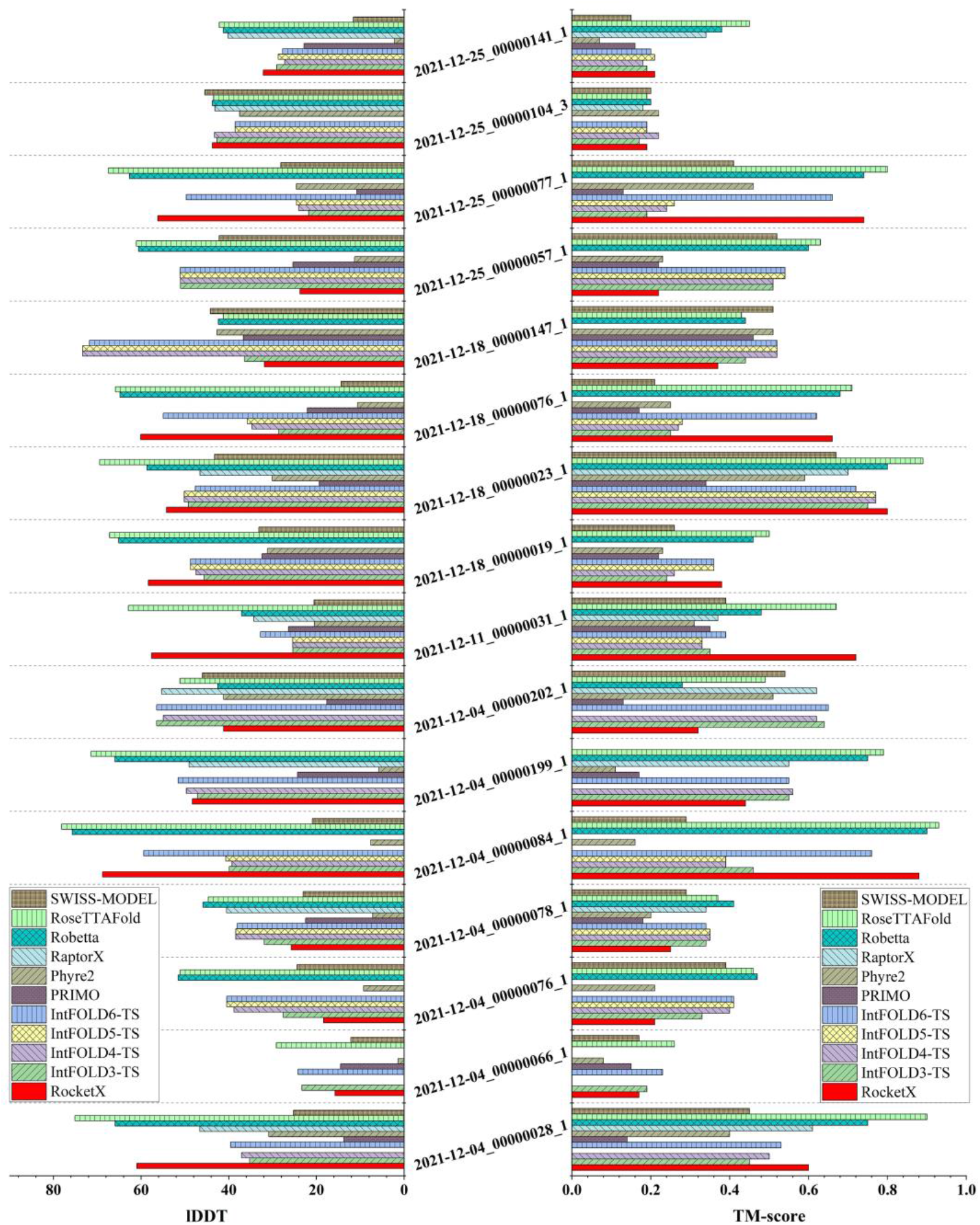
Performance of RocketX and the servers in CAMEO on the hard targets of 1 month blind test (2021-11-04 to 2021-12-25) of CAMEO. The left is the IDDT score, and the right is the TM-score.

RocketX achieves higher TM-score and lDDT than IntFold6-TS on 9 targets. In addition, RocketX has obtained models with TM-score≥0.8 on 2 targets, Robetta and RoseTTAFold have 2 and 4, respectively, and other methods have not. In particular, RocketX achieves the highest TM-score (0.72) on the target 2021-12-11_00000031_1, which is 7.46% higher than that (0.67) of RoseTTAFold. On the 29 medium targets, the average TM-score of RocketX is 0.73, which is lower than those of RoseTTAFold and IntFold6-TS but higher than those of other methods, ranking 3rd out of 11 methods. RocketX has obtained models with TM-score≥0.8 on 18 targets, which is 1 more than IntFold6-TS and 2 less than RoseTTAFold. In particular, RocketX achieves the highest TM-score (0.84) on the target 2021-12-04_00000033_3. Figure 7 shows the superimpositions of the model predicted by RocketX and the experimental structure of the CAMEO targets 2021-12-11_00000031_1, 2021-12-04_00000033_3.

**Fig. 7.**
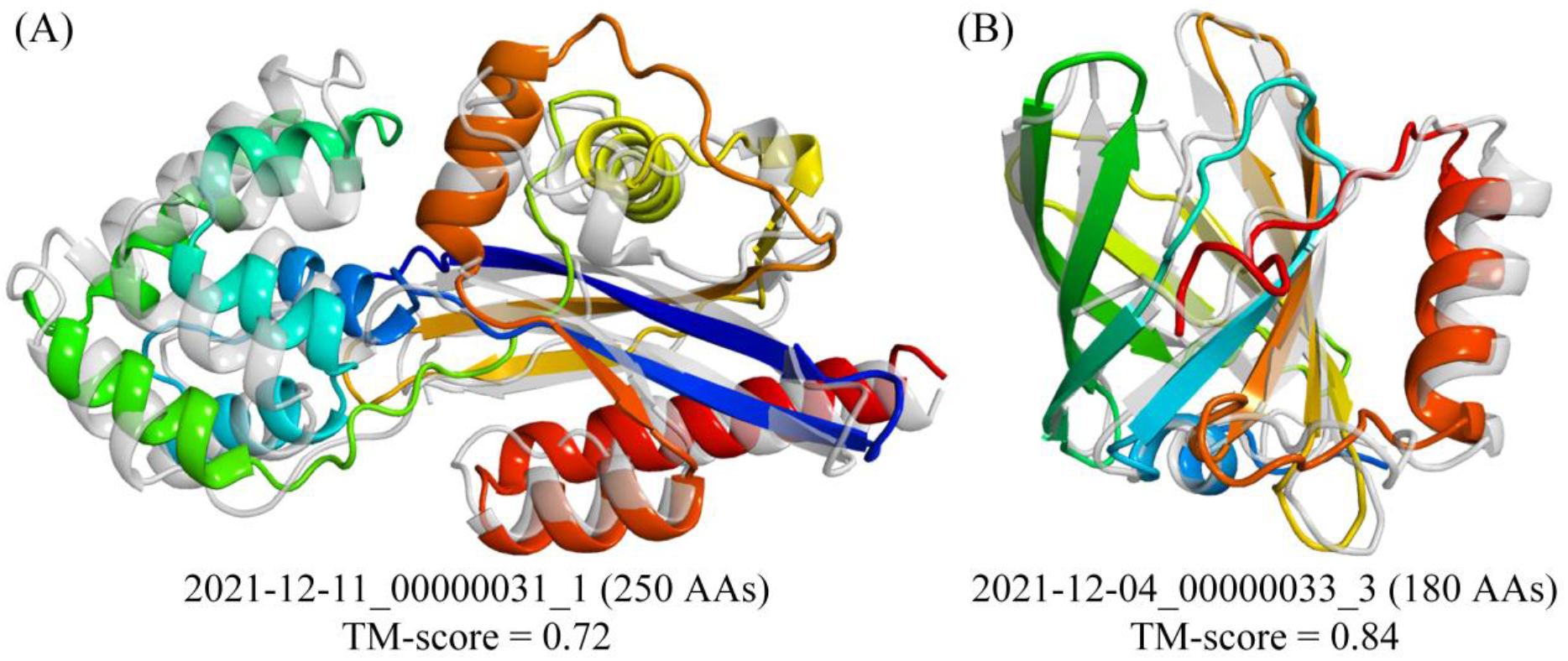
Superimposition between the model (rainbow) predicted by RocketX and the experimental structure (gray) of the CAMEO targets 2021-12-11_00000031_1, 2021-12-04_00000033_3.

## 4 Conclusion

We propose a de novo protein structure prediction method called RocketX by incremental inter-residue geometries prediction and model quality assessment using deep learning. In RocketX, a closed-loop feedback mechanism is constructed by inter-residue geometries prediction (GeomNet), structure simulation, and model quality assessment (EmaNet). In GeomNet, the MSA of the query sequence is first searched from the sequence databases, and the co-evolutionary features extracted from the MSA are fed into a deep residual network to predict the inter-residue geometric constraints. The structure model is generated through a simulation based on the predicted geometric constraints, which will be evaluated by EmaNet. In EmaNet, the features extracted from the structure model are sent to a deep residual network to estimate the inter-residue distance deviation and per-residue lDDT. The estimate results will be fed back to GeomNet as dynamic features to improve the inter-residue geometries prediction and guide structure folding. The experimental results show that our method outperforms the state-of-the-art methods trRosetta (without templates) and RaptorX; and that the model assessment feedback mechanism significantly contributes to the improved performance of RocketX. The blind test of CAMEO also shows that the performance of RocketX on medium and hard targets is comparable to the advanced methods that integrate templates even when no template is used in RocketX. With the rapid improvement in the accuracy of structure prediction, the protein folding paths and multiple states of protein structure will be investigated in the future.

